# H3K4 trimethylation is required for postnatal pancreatic endocrine cell functional maturation

**DOI:** 10.1101/2020.11.29.402990

**Authors:** Stephanie A. Campbell, Jocelyn Bégin, Cassandra L. McDonald, Ben Vanderkruk, Tabea L. Stephan, Brad G. Hoffman

## Abstract

During pancreas development, endocrine progenitors differentiate into the islet-cell subtypes, which undergo further functional maturation in postnatal islet development. In islet β-cells, genes involved in glucose-stimulated insulin secretion are activated and glucose exposure increases the insulin response as β-cells mature. Here, we investigated the role of H3K4 trimethylation in endocrine cell differentiation and functional maturation by disrupting TrxG complex histone methyltransferase activity in mouse endocrine progenitors. In the embryo, genetic inactivation of TrxG component *Dpy30* in NEUROG3+ cells did not affect the number of endocrine progenitors or endocrine cell differentiation. H3K4 trimethylation was progressively lost in postnatal islets and the mice displayed elevated random and fasting glycemia, as well as impaired glucose tolerance by postnatal day 24. Although postnatal endocrine cell proportions were equivalent to controls, islet RNA-sequencing revealed a downregulation of genes involved in glucose-stimulated insulin secretion and an upregulation of immature β-cell genes. Comparison of histone modification enrichment profiles in NEUROG3+ endocrine progenitors and mature islets suggested that genes downregulated by loss of H3K4 trimethylation more frequently acquire active histone modifications during maturation. Taken together, these findings suggest that H3K4 trimethylation is required for the activation of genes involved in the functional maturation of pancreatic islet endocrine cells.

## Introduction

Endocrine cell specification from pancreatic progenitors is initiated by the induction of the pro-endocrine factor *Neurog3* and subsequent exit from the cell cycle (Jensen et al., 2000; Larsen and Grapin-Botton, 2017; Miyatsuka et al., 2011). NEUROG3 drives expression of downstream transcription factors, such as *Neurod1, Nkx2-2, Nkx6-1, Pax6* and *Pdx1*, which determine further differentiation of endocrine progenitors into hormone-expressing endocrine cells that migrate and coalesce to form proto-islet structures (Bastidas-Ponce et al., 2017; Larsen and Grapin-Botton, 2017). For example, NEUROD1, NKX6-1 and PDX1 drive differentiation of insulin-secreting pancreatic β-cells (Bastidas-Ponce et al., 2017; Schaffer et al., 2013). In the first few weeks after birth, an increase in endocrine cell proliferation drives islet remodeling into the mature, spherical islet architecture (Bastidas-Ponce et al., 2017; Larsen and Grapin-Botton, 2017). Endocrine cells undergo functional maturation during this postnatal period which involves acquisition of glucose-sensing and hormone-secretion machinery (Bastidas-Ponce et al., 2017; Blum et al., 2012; Larsen and Grapin-Botton, 2017; Liu and Hebrok, 2017). For β-cell maturation in particular, the transition from immature to mature β-cells involves activation of the maturity marker *Ucn3* (Blum et al., 2012; van der Meulen et al., 2015), metabolic gene expression changes (e.g. switch from high-affinity hexokinase (*Hk*) to low-affinity glucokinase (*Gck*)) and improved glucose-stimulated insulin secretion due to increased glucose exposure (Liu and Hebrok, 2017; Stolovich-Rain et al., 2015). In addition, mature β-cells are defined by the repression of immature or “disallowed” β-cell genes that are often detected in stem cell-derived pancreatic β-like cells and in models of diabetes (Dhawan et al., 2015; Kieffer, 2016; Liu and Hebrok, 2017).

The gene expression changes that occur during endocrine cell differentiation and maturation are associated with specific histone modifications surrounding *cis*-regulatory loci. In addition to histone H3 lysine 27 acetylation (H3K27ac), genes that become activated during pancreas and endocrine cell differentiation are also associated with histone H3 lysine 4 monomethylation (H3K4me1) at enhancers and H3K4 trimethylation (H3K4me3) at promoters (Heintzman et al., 2009; 2007; Tennant et al., 2013). The majority of H3K4 methylation is catalyzed by the Trithorax Group (TrxG) complexes, which each contain one of six mammalian histone methyltransferases (i.e. SET1A/B, MLL1-4) and a minimum of four core proteins (i.e. ASH2L, DPY30, RBPP5 and WDR5) (Bochyńska et al., 2018; Campbell and Hoffman, 2016; Schuettengruber et al., 2017). Although H3K4 methylation is associated with active chromatin, several studies have suggested that H3K4 methylation is dispensable for gene expression (Dorighi et al., 2017; Hödl and Basler, 2012; Pérez-Lluch et al., 2015; Rickels et al., 2017; Xie et al., 2020).

Previously, we reported that loss of *Dpy30* and H3K4 methylation from PDX1^+^ progenitors increased the proportion of CPA1^+^ acinar progenitors while NEUROG3^+^ endocrine progenitors were reduced, suggesting a role for H3K4 methylation in endocrine cell specification (Campbell et al., 2019). While loss of H3K4 trimethylation had a minimal effect on bulk acinar or endocrine cell gene expression, our data suggested a role for H3K4 trimethylation in transcriptional maintenance of highly expressed genes in the acinar lineage. However, this model left unresolved whether H3K4 trimethylation is required for the transcription of genes essential for islet endocrine cell differentiation or functional maturation. As such, in this study, our objective was to determine the role of H3K4 trimethylation in the differentiation and functional maturation of mouse pancreatic endocrine cells.

## Results

### H3K4 trimethylation is lost postnatally in Dpy30ΔN islets

To investigate the role of H3K4 trimethylation in endocrine cell differentiation and maturation, we disrupted the TrxG complex core component *Dpy30* in NEUROG3^+^ endocrine progenitor cells using *Neurog3*-Cre driver mice (Schonhoff et al., 2004). DPY30 is required for TrxG catalytic activity or H3K4 methylation without affecting complex formation or co-activator activity (Bertero et al., 2015; Haddad et al., 2018). Previously described *Dpy30*^flox/flox^ mice (Campbell et al., 2019) were bred to generate experimental *Neurog3*-Cre; *Dpy30*^flox/flox^ mice (hereafter known as *Dpy30ΔN*) and littermate *Dpy30*^flox/flox^ (Cre-negative) control mice. In *Dpy30ΔN* embryos, DPY30 staining was absent from endocrine cells detected with the pan-endocrine marker chromogranin A (CHGA) beginning at embryonic day 14.5 (E14.5) and at postnatal day 0 (P0), P7, P14 and P24 (Figure 1A). However, DPY30 staining was maintained in the surrounding *Dpy30ΔN* exocrine pancreas, validating deletion of *Dpy30* in the endocrine lineage. At P24, Cre recombination was highly efficient in *Dpy30ΔN* islets as DPY30 staining was not detected in 95% of CHGA^+^ endocrine cells (Figure 1B). Co-staining of endocrine cells with CHGA and H3K4me3 revealed a delay between loss of DPY30 protein and H3K4 methylation (Figure 1C). Consistently, H3K4me3 staining was absent from 95% of CHGA^+^ *Dpy30ΔN* endocrine cells at P24 (Figure 1D). Measuring the median H3K4me3 intensity in CHGA^+^ islets relative to surrounding non-CHGA^+^ H3K4me3 intensity levels revealed that H3K4me3 intensity was maintained in E14.5 CHGA^+^ *Dpy30ΔN* endocrine cells similar to controls (Figure 1E). However, H3K4me3 intensity was progressively decreased from P0 onward and was consistently absent from P14 endocrine cells and thereafter (Figure 1E). At P24, Western blot analysis confirmed that both H3K4me3 (~4.5-fold, *P* < 0.01) and H3K4me1 (~2.2-fold, *P* < 0.05) were significantly reduced in *Dpy30ΔN* islets (Figure 1F). These data suggest that disruption of *Dpy30* in NEUROG3^+^ endocrine progenitors results in loss of DPY30 protein from embryonic endocrine cells and in the delayed loss of H3K4 trimethylation in postnatal islets.

**Figure 1:**
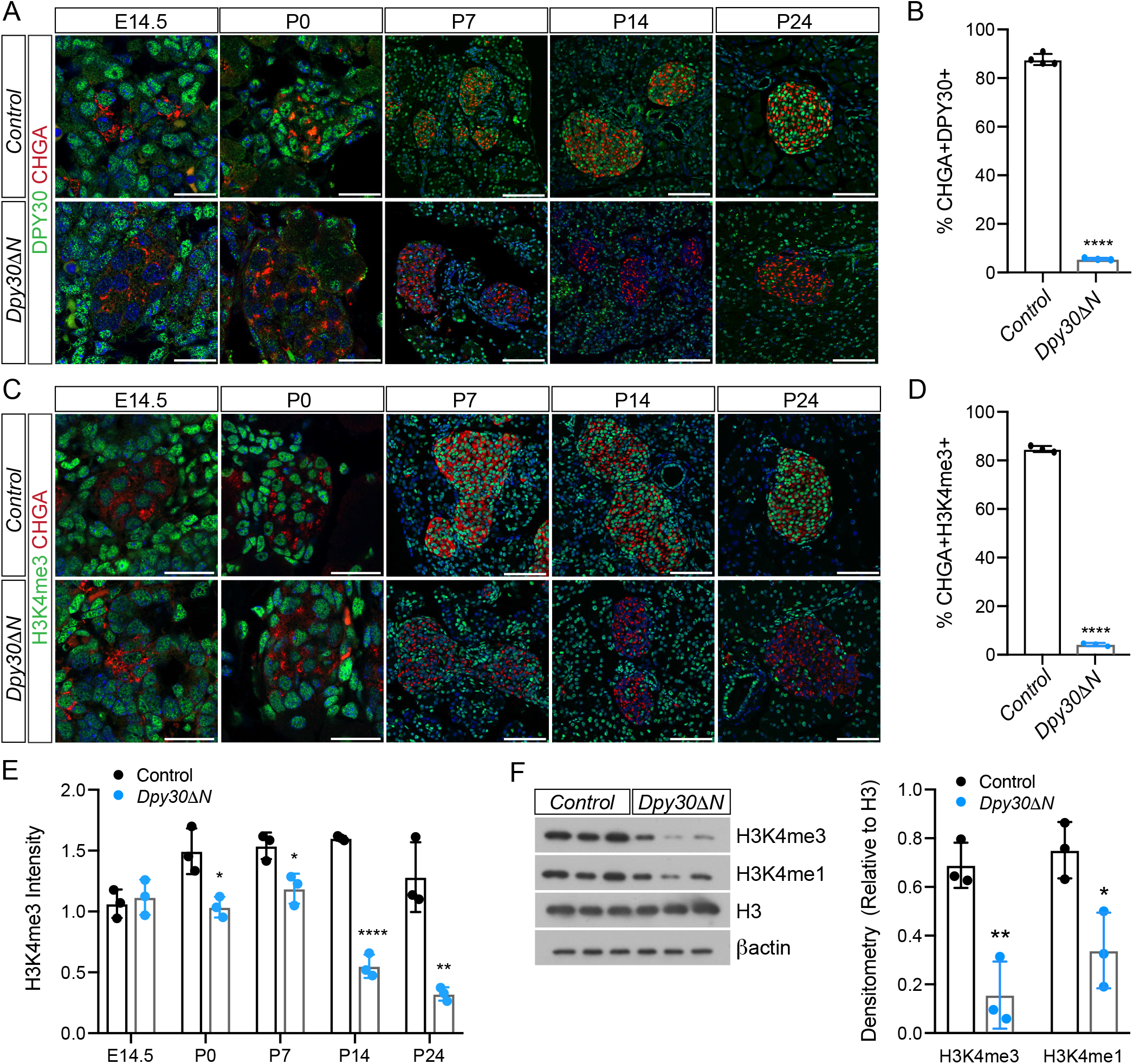
*Dpy30* deletion in pancreas endocrine progenitor cells leads to progressive loss of H3K4 methylation in *Dpy30*Δ*N* islets. **(A)** Staining of DPY30, chromogranin A (CHGA) and nuclei (blue) in control and *Dpy30*Δ*N* pancreas from E14.5 to P24. **(B)** The percentage of CHGA+DPY30+ cells in P24 control and *Dpy30*Δ *N* pancreas. **(C)** Staining of H3K4me3, CHGA and nuclei (blue) in control and *Dpy30*Δ*N* pancreas from E14.5 to P24. **(D)** The percentage of CHGA+/H3K4me3+ cells in P24 control and *Dpy30*Δ*N* pancreas. **(E)** The median H3K4me3 intensity in control and *Dpy30*Δ*N* CHGA+ cells relative to surrounding non-CHGA+ cell H3K4me3 intensity from E14.5 to P24. **(F)** Western blot and densitometry for H3K4me3 and H3K4me1 relative to histone H3 from P24 control and *Dpy30*Δ*N* islets. Data is presented as mean ± SD; n > 3; *Dpy30*Δ*N* vs. control; unpaired, two-tailed Student’s t-test. Scale bar, 25 μm (E14.5, P0) and 75 μm (P7-P24).

### Loss of Dpy30 does not affect NEUROG3^+^ endocrine progenitor cell specification

To determine whether deletion of *Dpy30* in NEUROG3^+^ endocrine progenitors altered the proportion of islet endocrine cells, we first examined the ratio of NEUROG3^+^ endocrine progenitors to SOX9^+^ pancreas progenitors and determined there was no significant difference in E14.5 control and *Dpy30ΔN* pancreas (Figure 2A-B). Next, co-staining for insulin^+^ β-cells, glucagon^+^ α-cells and somatostatin^+^ δ-cells demonstrated that the sum of these endocrine cell types relative to total pancreas cells was not altered in the E18.5 *Dpy30ΔN* pancreas compared to controls (Figure 2C-D). The relative proportion of *Dpy30ΔN* islet insulin^+^ β-cells, glucagon^+^ α-cells and somatostatin^+^ δ-cells was also unaffected both in E18.5 pancreas (Figure 2E) and postnatally in P24 *Dpy30ΔN* pancreas (Figure 2F-G). Although there were no changes in islet endocrine cell proportions, we noted that the insulin staining in *Dpy30ΔN* islets appeared reduced compared to controls (Figure 2F). Further, quantification of CHGA^+^ area at P24 revealed no changes in total endocrine cell area between the control and *Dpy30ΔN* pancreas (Figure 2H). These data suggest that disruption of *Dpy30* in NEUROG3^+^ progenitors does not affect the proportion of differentiated endocrine cells in the embryo or postnatal pancreas.

**Figure 2:**
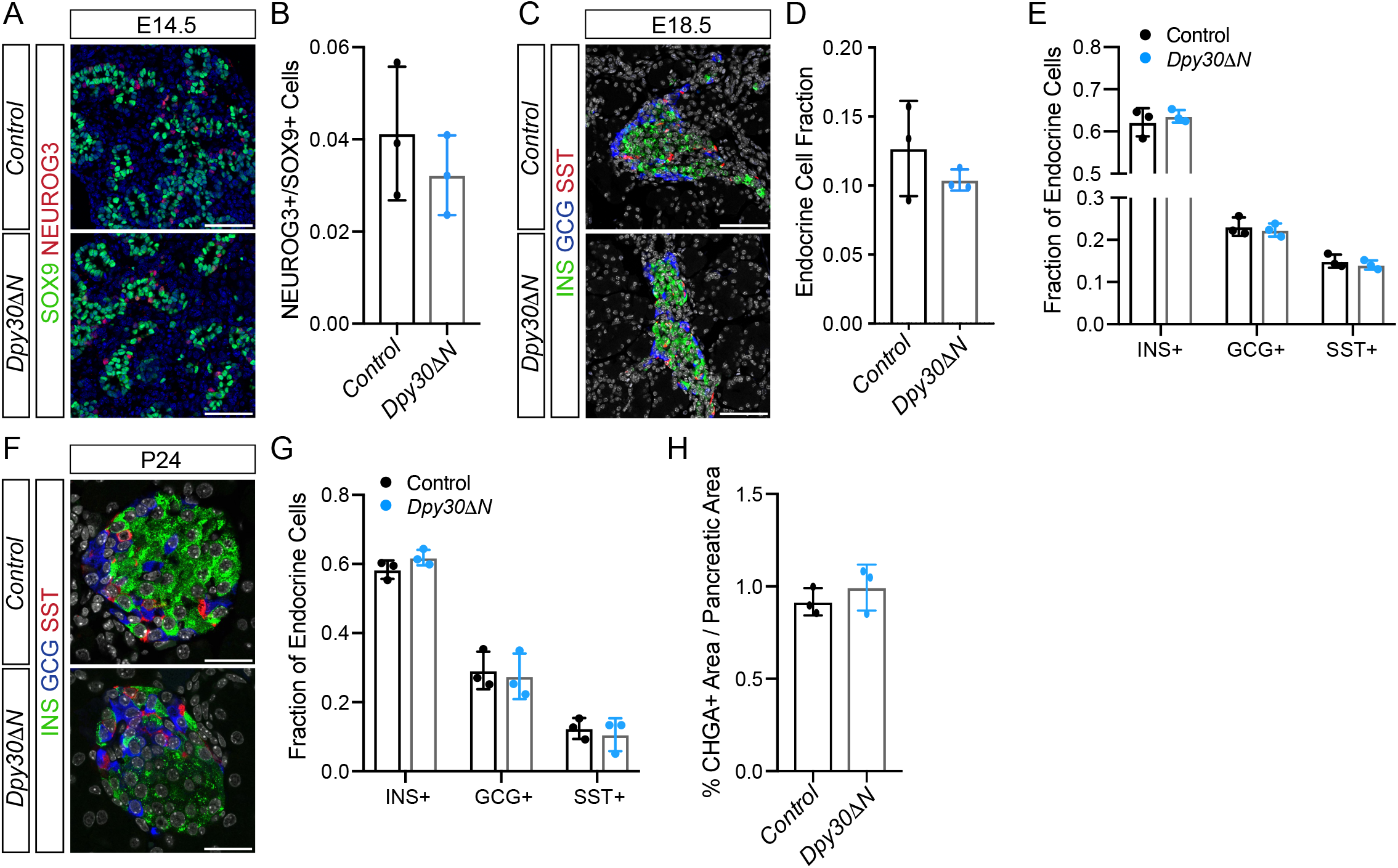
The proportion of endocrine cells is not altered in *Dpy30*Δ*N* mice. **(A)** Staining for SOX9, NEUROG3 and nuclei (blue) in E14.5 control and *Dpy30*Δ*N* pancreas. **(B)** The NEUROG3+/SOX9+ cell ratio in E14.5 control and *Dpy30*Δ*N* pancreas. **(C)** Co-staining for insulin (INS), glucagon (GCG), somatostatin (SST) and nuclei (grey) in E18.5 control and *Dpy30*Δ*N* pancreas. **(D)** The fraction of endocrine cells (sum of INS+, GCG+ and SST+ cells) relative to pancreas cells in E18.5 control and *Dpy30*Δ*N* pancreas. **(E)** The proportions of insulin+ β-cells (INS+), glucagon+ α-cells (GCG+) and somatostatin+ δ-cells (SST+) relative to total endocrine cells in E18.5 control and *Dpy30*Δ*N* pancreas. **(F)** Co-staining for insulin (INS), glucagon (GCG), somatostatin (SST) and nuclei (grey) in P24 control and *Dpy30*Δ*N* pancreas. **(G)** The proportions of INS+, GCG+ and SST+ cells relative to total endocrine cells in P24 control and *Dpy30*Δ*N* pancreas. **(H)** Quantification of CHGA+ area in P24 control and *Dpy30*Δ*N* pancreas. Data are presented as mean ± SD; n = 3; *Dpy30*Δ*N* vs. control; unpaired, two-tailed Student’s t-test. Scale bar, 75 μm.

### Dpy30ΔN mice develop hyperglycemia and impaired glucose tolerance

To investigate whether the postnatal islet reduction in H3K4 methylation had a biological effect in *Dpy30ΔN* mice, we first monitored body mass and random fed blood glucose levels. Daily tracking between P24 and P38 indicated that while body mass was not altered in male *Dpy30ΔN* mice compared to controls (Figure 3A), mean *ad libitum* glycemia was over 18 mM by P28 and worsened over time (Figure 3B). A similar trend was observed in female *Dpy30ΔN* mice (Figure S1A-B). Following a 6 hour fast at P23, blood glucose levels were equivalent in control and *Dpy30ΔN* male mice (Figure S2A). Intraperitoneal glucose tolerance tests (IPGTTs) at this stage (P23) revealed a marginal increase to ~19 mM at 30 minutes, although blood glucose measurements at other time points were otherwise comparable to controls (Figure S2B). At P24, although fasting blood glucose levels remained similar between control and *Dpy30ΔN* male mice (Figure 3C), blood glucose was significantly elevated in *Dpy30ΔN* males following an IP glucose challenge (Figure 3D). At P25, fasting glycemia levels were elevated to ~12 mM in *Dpy30ΔN* male mice (Figure S2C) and blood glucose levels were above the limit of detection (33.3 mM) at 15- and 30-minutes post-glucose injection (Figure S2D). Finally, we measured islet insulin content and detected a ~40% reduction in insulin levels from P24 male *Dpy30ΔN* islets relative to controls (Figure 3E). A ~50% reduction in islet insulin content was also detected in female *Dpy30ΔN* mice (Figure S1C). These data suggest that *Dpy30ΔN* mice develop random and fasting hyperglycemia, impaired glucose tolerance and have decreased islet insulin content.

**Figure 3:**
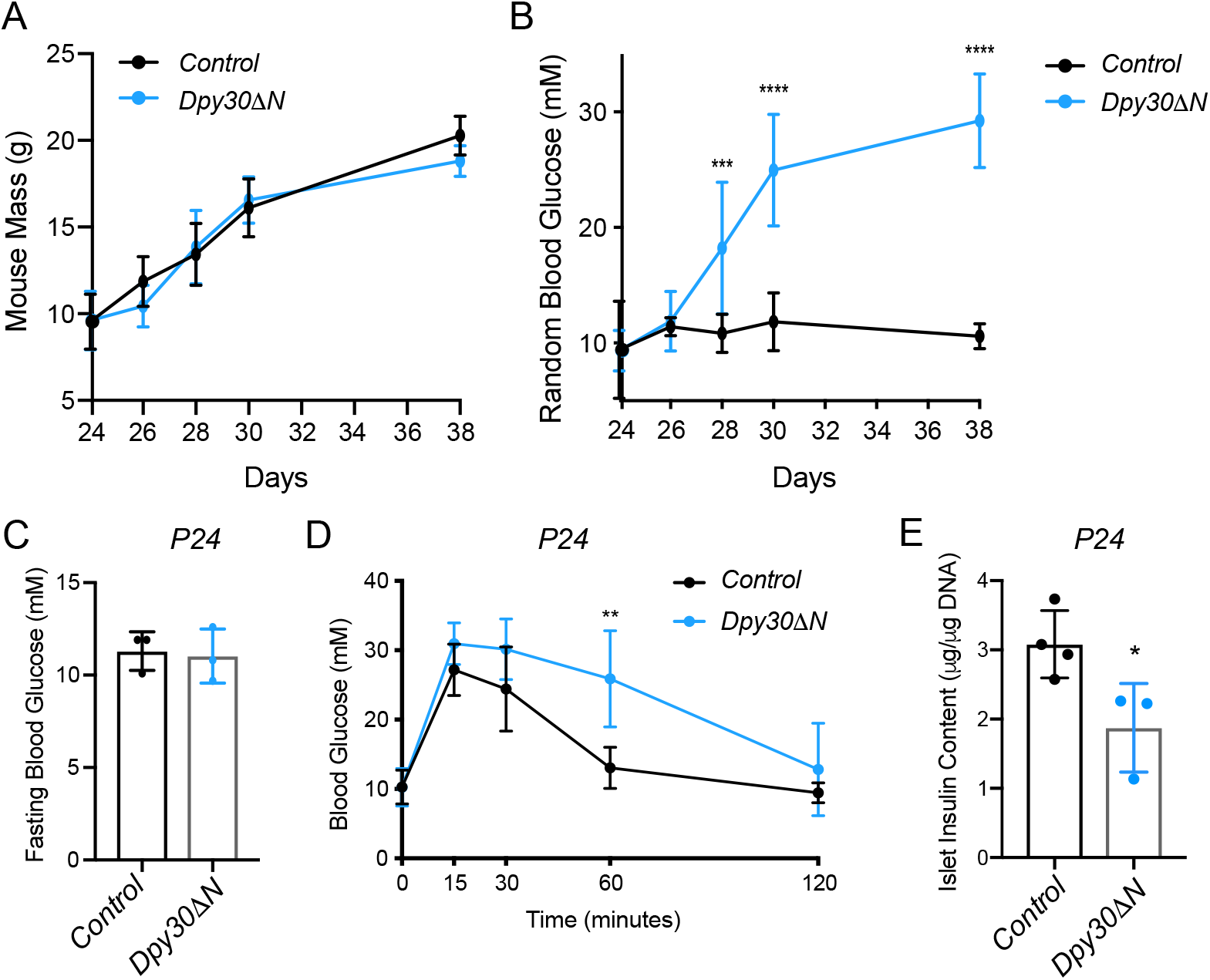
*Dpy30*Δ*N* mice develop hyperglycemia and impaired glucose tolerance. **(A)** Mouse body mass measurements between P24 and P38 in male control and *Dpy30*Δ*N* animals. **(B)** Random blood glucose measurements from male *Dpy30*Δ*N* and control mice between P24 and P38. **(C)** Blood glucose measurements after a 6 hour fast in P24 male control and *Dpy30*Δ*N* mice. **(D)** IPGTT of 2 g per kg body mass IP glucose in P24 male control and *Dpy30*Δ*N* mice following a 6 hour fast. **(E)** Islet insulin content relative to DNA from P24 male control and *Dpy30*Δ*N* islets. Data are presented as mean ± SD; n > 3; *Dpy30*Δ*N* vs. control; multiple t tests with Holm-Sidak correction for multiple comparisons (in A and B); unpaired, two-tailed Student’s t test (in C and E); repeated measures two-way ANOVA with Sidak’s multiple comparison post-hoc test (in D). Note that statistical testing was not performed at 15 and 30 minutes (in D) as measurements were above the detection limit (33.3 mM).

### H3K4 trimethylation is necessary for the expression of mature pancreatic islet genes

To understand the underlying mechanism for hyperglycemia and impaired glucose tolerance in *Dpy30ΔN* mice, we next examined whether islet dysfunction was the result of reduced endocrine cell maturity. To assess islet maturation, islets from male *Dpy30ΔN* and control mice were collected at P24 and RNA-sequencing (RNA-seq) was performed. Differential gene expression analysis demonstrated that endocrine cell hormones (e.g. *Gcg, Ins1, Ins2, Ppy* and *Sst*) and genes involved in insulin secretion or β-cell maturation (e.g. *Abcc8, Glp1r, Kcnj11, Slc2a2, Slc30a8* and *Ucn3*) were downregulated in *Dpy30ΔN* islets compared to controls (Figure 4A-B). Interestingly, the glucagon receptor (*Gcgr*) was one of the most downregulated genes (~16-fold reduction) in *Dpy30ΔN* islets. Genes in the glucose-stimulated insulin secretion pathway were particularly affected by decreased H3K4 methylation in maturing pancreatic islets (Figure 4C). Elevated genes in *Dpy30ΔN* islets included genes associated with immature β-cells (e.g. *Aldh1a3, Gast, Hk1/2, Ldha, Npy, Pdgfra, Rest and Slc16a1*) and several genes involved in cell adhesion (e.g. *Apoa1, Ccn3, Cldn2, Col16a1, Itga11* and *Lamc3*) (Figure 4A-B). RNA-seq analysis also revealed that neither mature islet transcription factors (e.g. *Mafa, Neurod1, Nkx2-2, Nkx6-1, Pax6* and *Pdx1*) nor genes involved in mTOR signaling (e.g. *Akt1, Mtor, Pdk1* and *Rptor*) were differentially expressed in *Dpy30ΔN* islets (Figure S3). GO term analysis indicated that genes involved in ion and transmembrane transport and hormone and insulin secretion were reduced in *Dpy30ΔN* cells, while genes involved in extracellular matrix formation and cell adhesion were elevated (Figure 4D).

**Figure 4:**
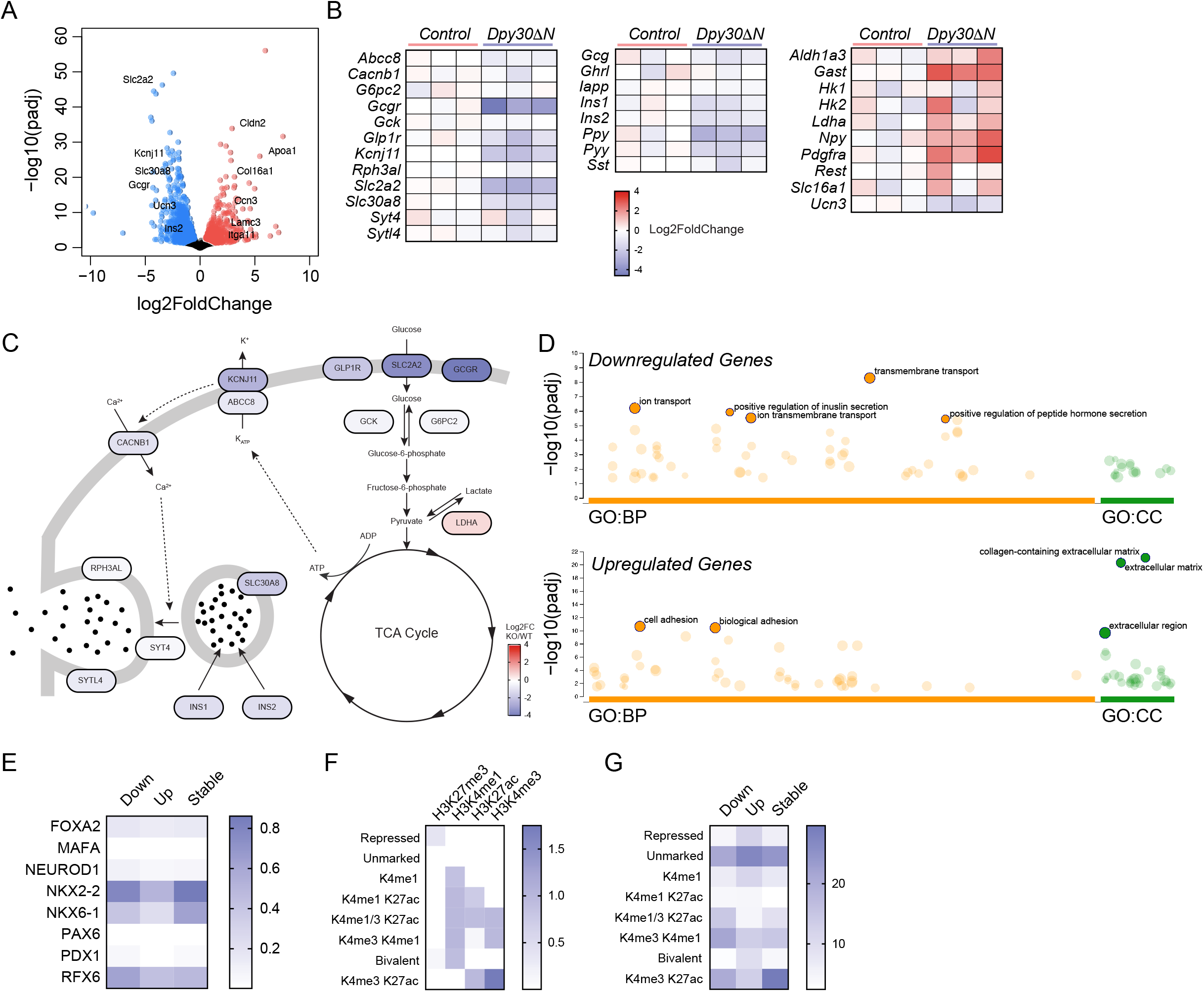
Islet maturity is compromised in *Dpy30*Δ*N* mice. **(A)** Volcano plot of differentially expressed genes in P24 *Dpy30*Δ*N* vs. control islets with log2 fold change plotted against the adjusted p-value (−log10). Downregulated genes are in blue, upregulated genes are in red and stable genes are in black. **(B)** Heatmaps of select gene expression (log2 fold change) comparing 3 biological replicates across P24 control and *Dpy30*Δ*N* islets. **(C)** Overview of the glucose-stimulated insulin secretion pathway with affected genes in P24 *Dpy30*Δ*N* islets represented by log2 fold change values. **(D)** g:Profiler GO term analysis of downregulated and upregulated genes from P24 control and *Dpy30*Δ*N* islets represented as adjusted p-values (−log10). BP = biological process; CC = cellular component. **(E)** Heatmap of transcription factor occupancy from adult islet ChIP-seq data at downregulated, upregulated and stable genes in P24 *Dpy30*Δ*N* vs. control islets. **(F)** Histone modification enrichment plot of adult islet ChIP-seq data indicating repressed (H3K27me3), unmarked, poised (H3K4me1/3), active (H3K4me1/3 and H3K27ac) and bivalent (H3K4me1 and H3K27me3) chromatin states. **(G)** Heatmap showing the fraction of downregulated, upregulated and stable genes in P24 *Dpy30*Δ*N* vs. control islets in each chromatin state identified in **(F)**.

Transcription factor occupancy and histone modification status at downregulated, upregulated and stably expressed genes in *Dpy30ΔN* islets was examined using previously published adult mouse islet chromatin immunoprecipitation sequencing (ChIP-seq) data (Gutiérrez et al., 2017; Lu et al., 2018; Piccand et al., 2014; Swisa et al., 2017; Taylor et al., 2013; Tennant et al., 2013). The islet transcription factors NKX2-2, NKX6-1 and RFX6 were marginally depleted at upregulated genes (Figure 4E). We utilized ChromHMM to stratify the histone modification ChIP-seq data into chromatin states, including repressed (H3K27me3), unmarked, poised (H3K4me1/3), active (H3K4me1/3 and H3K27ac) and bivalent (H3K4me1 and H3K27me3) chromatin (Figure 4F). Histone modifications associated with active chromatin were relatively increased at the promoters of downregulated and stable genes (Figure 4G). In contrast, the *cis*-regulatory elements governing the upregulated genes were in an inactive chromatin state, with either unmarked histones or enrichment of repressive, bivalent or poised histone modifications (Figure 4G). Together, these results suggest that hyperglycemia and impaired glucose tolerance in *Dpy30ΔN* mice manifests from the reduced expression of genes required for islet β-cell maturity and glucose-stimulated insulin secretion.

### Failed activation of pro-endocrine cis-regulatory elements in Dpy30ΔN mice

To explore whether the differentially expressed genes in P24 *Dpy30ΔN* islets require distinct chromatin state changes during pancreatic endocrine cell development to become active, we assessed the histone modification profiles of the genes in *Neurog3*^HI^ cells compared to adult islets (Gutiérrez et al., 2017; Piccand et al., 2014; Swisa et al., 2017; Taylor et al., 2013; Tennant et al., 2013; Yu et al., 2018). Our analysis revealed that in *Neurog3*^HI^ endocrine progenitors downregulated, upregulated and stable genes had a similar H3K4me3 profile at their promoters; however, there was noticeably increased enrichment of H3K4me3 at downregulated in islets as compared to in *Neurog3*^HI^ endocrine progenitors (Figure 5A). The H3K4me1 profiles revealed that promoters of downregulated and upregulated genes were blocked in *Neurog3*^HI^ cells (Cheng et al., 2014), whereas H3K4me1-marked histones shifted away from the transcription start site (TSS) in adult islets (Figure 5B). H3K27ac enrichment profiles were relatively unchanged between *Neurog3*^HI^ cells and adult islets, more than for upregulated genes (Figure 5C). H3K27me3 profiles were also similar for downregulated, upregulated and stable genes in *Neurog3*^HI^ cells, although upregulated genes had modestly increased enrichment of the repressive H3K27me3 histone modification in adult islets (Figure 5D). Assessment of transitions from the chromatin states identified in *Neurog3*^HI^ endocrine progenitors to those identified in mature endocrine islet cells showed that an increased fraction of genes that were downregulated in *Dpy30ΔN* islets transitioned to an active chromatin state (H3K27ac and/or H3K4me1/3) during development (Figure 5E-F). Conversely, genes that were upregulated in *Dpy30ΔN* islets more frequently gained a repressive chromatin state (H3K27me3, bivalent or unmarked) in adult islets (Figure 5E-F). UCSC Genome Browser views at the TSS and *cis*-regulatory loci of *Glp1r, Slc2a2* and *Slc30a8* demonstrate acquisition of active histone marks during endocrine cell maturation at these loci (Figure 5G). Overall, these data suggest there is a marginal increase in the fraction of downregulated genes in *Dpy30ΔN* islets that acquire active histone modifications during development, while upregulated genes are more likely to acquire a repressive state.

**Figure 5:**
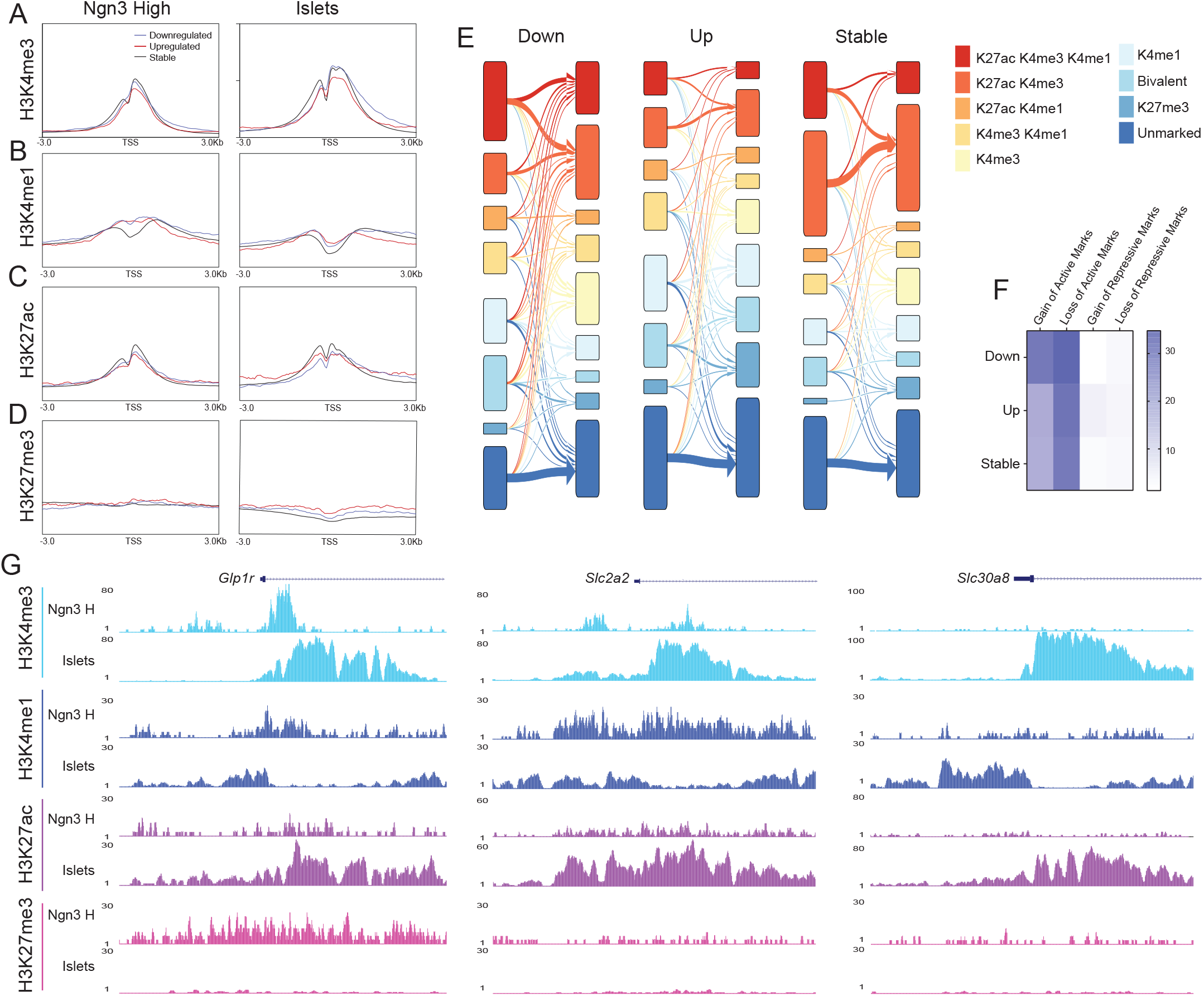
Genes downregulated in *Dpy30*Δ*N* mice require activation during endocrine cell differentation. **(A-D)** Comparison of histone modification profiles in E13.5 *Neurog3*^*HI*^ cells vs. adult islets around the TSS of downregulated, upregulated and stable genes in P24 *Dpy30*Δ*N* vs. control islets. Distributions for **(A)**H3K4me3, **(B)** H3K4me1, **(C)** H3K27ac and **(D)** H3K27me3 are shown ± 3.0 kb around the TSS. **(E)** Alluvial plot of chromatin state transitions for P24 islet *Dpy30*Δ*N* downregulated, upregulated and stable genes from *Neurog3*^*HI*^ cells to islets. **(F)** Heatmap summarizing the overall chromatin state transitions for P24 islet *Dpy30*Δ*N* downregulated, upregulated and stable genes from *Neurog3*^*HI*^ cells to islets. **(G)** Comparison of UCSC genome browser tracks at *Glp1r*, *Slc2a2* and *Slc30a8* for H3K4me3, H3K4me1, H3K27ac and H3K37me3 in *Neurog3*^*HI*^ cells and islets.

### H3K4 trimethylation is required for pancreatic endocrine cell maturation

To determine whether the endocrine cell terminal markers *Ins1, Ins2, Gcg, Sst* and *Ppy* that require activation during endocrine development fail to fully activate or instead lose expression in *Dpy30ΔN* islets, we used qPCR to quantify their expression at P7, P14 and P24. We found that expression of these markers at P7 was very similar between *Dpy30ΔN* and control islets (Figure 6A). The expression of all of these markers significantly increased in control islets by P24, but no increase was detected in *Dpy30ΔN* islets at P14 or P24 as compared to at P7 (Figure 6A). In contrast, *Npy* and *Gast*, which are known to have elevated expression in immature islets, were significantly increased at P24 in *Dpy30ΔN* islets as compared in control islets (Figure 6A). Staining pancreas sections from P24 control and *Dpy30ΔN* mice confirmed that gastrin protein levels were also elevated in *Dpy30ΔN* islets (Figure 6B). We next compared the expression of downregulated, upregulated and stable genes identified in our P24 islet RNA-seq analysis across endocrine cell differentiation and maturation. For this, we used previously published data from E13.5 *Neurog3*^HI^ endocrine progenitors and E17.5 INS1+ β-cells (Yu et al., 2018), as well as INS1+ β-cells from P1, P7, P14, P21 and P28 islets (Zeng et al., 2017). In agreement with our previous data, we found that an increased fraction of downregulated genes in *Dpy30ΔN* islets are normally activated during pancreatic β-cell maturation (Figure 6C). Combined, these data suggest that H3K4 trimethylation is necessary for the appropriate activation of mature endocrine cell genes during postnatal islet functional maturation.

**Figure 6:**
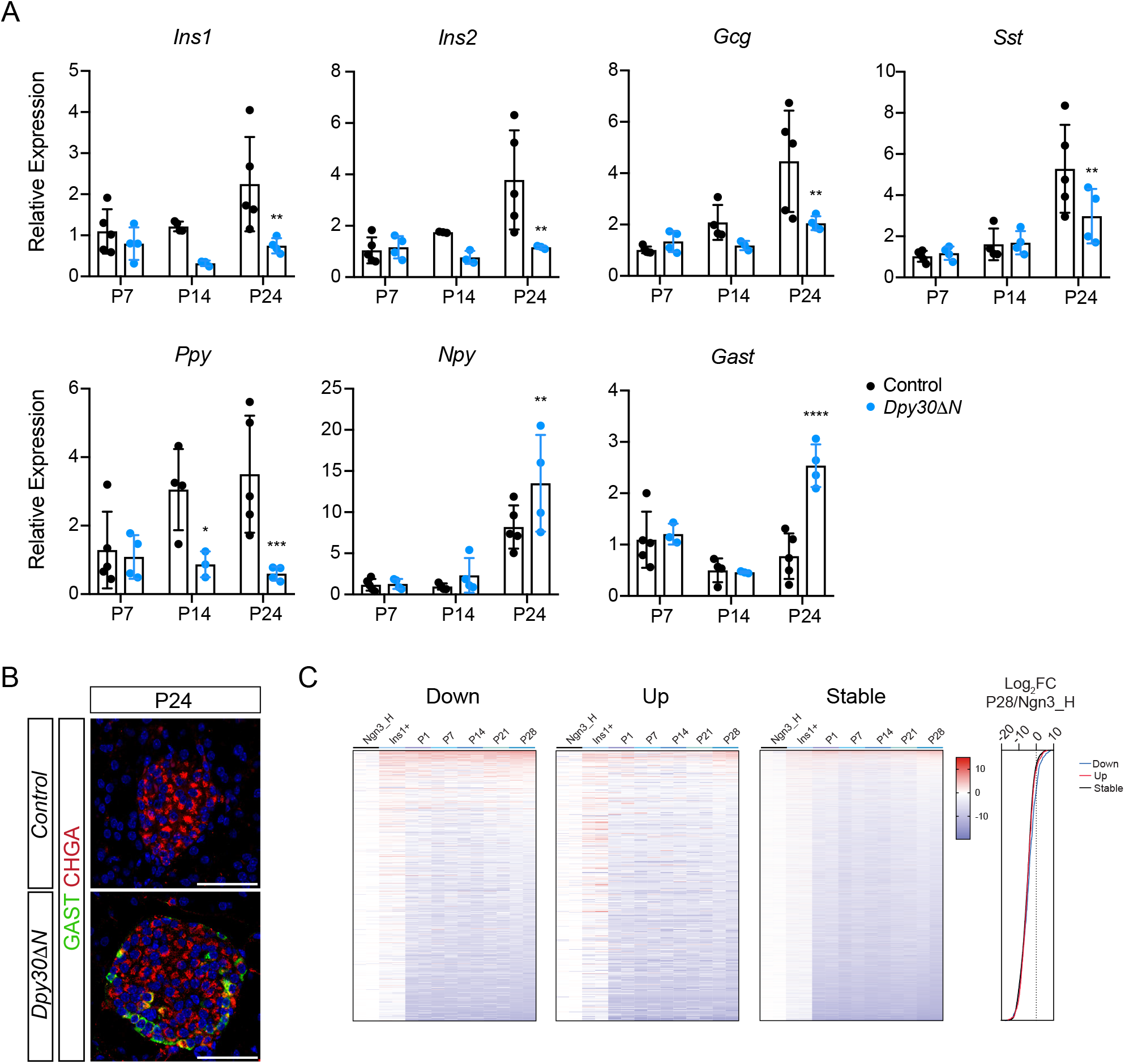
*Dpy30*Δ*N* endocrine islet cells do not undergo functional maturation. **(A)** Relative expression of *Ins1*, *Ins2*, *Gcg*, *Sst*, *Ppy*, *Npy* and *Gast* transcripts in P7, P14 and P24 control and *Dpy30*Δ*N* islets (n = 4-5) as determined by qPCR. **(B)** Staining for gastrin (GAST), CHGA and nuclei (blue) in P24 control and *Dpy30*Δ*N* pancreas. **(C)** Heatmaps of downregulated, upregulated and stable genes (identified in Figure 4) plotted as a developmental gene expression timeline. Data includes E13.5 5 *Neurog3*^*HI*^ endocrine progenitors, E17.5 INS1+ β-cells and β-cells from P1, P7, P14, P21 and P28 islets. Gene expression (log2foldchange) from P28 islets was compared to E13.5 *Neurog3*^*HI*^ endocrine progenitors for the downregulated, upregulated and stable gene sets. Data are presented as mean ± SD; *Dpy30*Δ*N* vs. control; unpaired, two-tailed Student’s t-test. Scale bar, 50 μm.

## Discussion

TrxG complex proteins are enriched in mature islets (Tennant et al., 2015) and interact with the maturation factor MAFA in mature β-cells (Scoville et al., 2015), suggesting a role for TrxG complexes and/or H3K4 methylation in maintenance of islet identity. Deletion of the TrxG gene *Ncoa6* in embryonic β-cells does not affect *Mafa* expression but results in reduced downstream activation of MAFA target genes and impaired glucose-stimulated insulin secretion (Scoville et al., 2015). Importantly, decreased gene expression was correlated with reductions in H3K4 trimethylation and RNA polymerase II (Pol II) at the TSS of affected genes. In addition, H3K4 trimethylation deposited by the SET7/9 histone methyltransferase was linked to transcriptional maintenance of genes involved in glucose-stimulated insulin secretion in primary islets (Deering et al., 2009). Together, these studies suggest that H3K4 trimethylation may play a role in maintaining β-cell gene expression, but whether H3K4 trimethylation is required during the process of endocrine cell maturation has not been examined.

In this study, we demonstrate that genetic inactivation of *Dpy30* in NEUROG3^+^ cells does not alter the proportion of pancreatic endocrine progenitors or endocrine cells. Global loss of H3K4 trimethylation occurred postnatally during islet-cell maturation. By P28, *Dpy30ΔN* mice displayed elevated random and fasting glycemia, in addition to impaired glucose tolerance by P24. These phenotypes could not be explained by changes in *Dpy30ΔN* endocrine cell area or the relative proportions of islet cells. However, islet insulin content and expression of insulin and genes associated with insulin secretion and β-cell maturity were reduced in P24 *Dpy30ΔN* islets. In addition, we detected impaired activation of *Ins1* and *Ins2* during the postnatal islet maturation period. Together, these data suggest that H3K4 trimethylation is required for normal islet-cell functional maturation.

### Hyperglycemia and impaired glucose tolerance correlates with loss of islet H3K4 trimethylation

As discussed above, loss of endocrine cell H3K4 mono- and trimethylation was delayed in *Dpy30ΔN* mice. Although DPY30 was absent from CHGA^+^ endocrine cells at E14.5, loss of H3K4me3 did not occur until P14. This difference is likely, at least in part, the result of failed H3K4me3 re-establishment during cell division. At the time when *Dpy30* is genetically inactivated, NEUROG3^+^ cells exit the cell cycle and differentiate into endocrine cells (Miyatsuka et al., 2011). These cells remain non-proliferative until the early postnatal period when endocrine cell replication increases (Cleaver and MacDonald, 2010), suggesting that there is progressive loss of H3K4 methylation after each round of endocrine cell division. Of note, we observed no phenotype in *Dpy30ΔN* mice prior to the loss of H3K4 methylation, suggesting that loss of DPY30 itself has a minimal effect and that the impaired endocrine cell maturation in *Dpy30ΔN* mice is due to the absence of H3K4 methylation.

The loss of H3K4 methylation from *Dpy30ΔN* islets at P14 overlaps with the period of endocrine cell maturation. A key determinant of β-cell maturity is the ability to effectively respond to glucose, which includes efficient glucose transport and metabolism as well as regulated insulin secretion. During the first few weeks after birth, endocrine cells undergo functional maturation and develop glucose-stimulated hormone secretion by P14 (Blum et al., 2012; Liu and Hebrok, 2017; Pan and Wright, 2011). Further β-cell maturation is triggered by exposure to a carbohydrate-rich diet after weaning at P21 (Stolovich-Rain et al., 2015). The transition to glucose metabolism as the primary energy source exposes maturing islets to increased glucose levels, and this stimulates β-cell proliferation, enhances insulin secretion and promotes oxidative phosphorylation (Stolovich-Rain et al., 2015). Following weaning, *Dpy30ΔN* mice developed hyperglycemia and impaired glucose tolerance beginning at P24, suggesting that either glucose-stimulated insulin secretion, insulin levels or insulin action may be affected at this stage. Measurement of islet insulin content and our gene expression data confirmed that insulin levels were significantly reduced in P24 *Dpy30ΔN* islets. In addition, we detected no significant difference in P24 islet area relative to controls, suggesting that hyperglycemia did not result from impaired β-cell proliferation. Instead, our data suggests that during a glucose challenge, *Dpy30ΔN* mice have insufficient insulin secretion to reduce blood glucose due to reduced levels of insulin.

### Pancreatic endocrine cell maturation is impaired in Dpy30ΔN mice

Specification of the pancreatic endocrine lineage begins with activation of *Neurog3* in a subset of SOX9+ pancreas progenitors in the developing embryo. NEUROG3+ cells delaminate and migrate away from the trunk epithelium into the surrounding mesenchyme, where they eventually differentiate into endocrine cells and coalesce into mature spherical islet structures (Bastidas-Ponce et al., 2017). Extracellular matrix proteins such as cadherins, collagens, integrins and laminins have an essential role in cell migration and cell-cell adhesion during endocrine cell aggregation into islet clusters (Bastidas-Ponce et al., 2017; Dahl et al., 1996; Stendahl et al., 2009). Interestingly, we detected a disproportionate upregulation of many genes associated with the extracellular matrix and cell-cell adhesion, including *Cldn2, Apoa1, Col16a1, Ccn3, Lamc3* and *Itga11* in P24 *Dpy30ΔN* islets, that play important roles in endocrine cell migration, isletogenesis, and maintenance of intercellular communication (Murtaugh et al., 2017).

The transition from immature to mature β-cells involves important gene expression changes, where genes exclusive to immature β-cells (e.g. *Hk1, Ldha, Rest and Pdgfra*) become repressed, and mature genes associated with insulin secretion machinery (e.g. *Gck, Slc2a2 and Slc30a8*) are induced (Liu and Hebrok, 2017). In *Dpy30ΔN* islets, the expression of several mature genes required for glucose transport (*Slc2a2*), glucose sensing (*Gck*) and insulin secretion (*Abcc8, Kcnj11*) was significantly reduced, in addition to the mature β-cell marker *Ucn3* (Blum et al., 2012). Meanwhile, several genes associated with endocrine cell immaturity, such as *Aldh1a3, Gast, Hk2, Pdgfra* and *Rest* were elevated in *Dpy30ΔN* islets. In addition, we detected GAST+ endocrine cells exclusively in P24 *Dpy30ΔN* islets. Furthermore, transcript levels of islet hormones (e.g. *Ins1*, *Ins2, Gcg, Ppy* and *Sst*) were also significantly reduced in *Dpy30ΔN* islets. Together, these data strongly suggest that at P24 *Dpy30ΔN* islet cells are still in an immature-like state. It is important to note that this immaturity is without significant changes to the expression of key islet transcription factors (e.g. *Mafa, Neurod1, Nkx2-2, Nkx6-1, Pax6* and *Pdx1*) (Figure S3). We also did not observe significant gene expression changes in the mTOR pathway in *Dpy30ΔN* islets (Figure S3), which plays an important role in islet functional maturation (Ni et al., 2017; Sinagoga et al., 2017).

### H3K4 methylation is acquired at mature β-cell genes during functional maturation

We found that genes upregulated in P24 *Dpy30ΔN* islets as compared were slightly more likely to become H3K27me3 marked or “unmarked” in islets as compared to in *Neurog3*^HI^ cells. This agrees with our gene expression data and suggests a subset of these genes are genes that are normally more abundant in progenitors or immature endocrine cells and that are generally “turned-off” in mature pancreas endocrine cells. It is less clear why some genes, specifically endocrine cell terminal markers such as *Ins1, Ins2, Gcg, Sst* and *Ppy*, and genes with functional relevance in β-cells fail to be activated in *Dpy30ΔN* endocrine cells, while other genes are unaffected. Our analysis of the chromatin state changes at these genes from *Neurog3*^HI^ to adult islets suggest they are more likely to be activated and to gain active chromatin marks, but otherwise are very similar in chromatin state to genes unaffected by the loss of H3K4me3. Other groups have found similar results, and have found that H3K4me3 loss primarily affects the expression of lineage-specific genes (Benayoun et al., 2014; Chen et al., 2015). Why this is the case, and whether H3K4 trimethylation is required purely to activate such genes, or if it is also essential to maintain lineage-specific expression is unknown and warrants further investigation.

Regardless, the results in this study suggest that H3K4 trimethylation is required for the expression of pancreatic islet genes involved in endocrine cell functional maturation. This conclusion is supported by evidence that H3K4 methylation is established at these genes during the endocrine cell maturation period. Overall, our data suggests that islet endocrine cells do not completely mature in the absence of H3K4 methylation.

## Acknowledgements

The authors would like to thank the staff of the Animal Care Facility at BCCHRI for daily maintenance of the mouse colonies. Funding was provided by British Columbia Children’s Hospital Research Institute, and the Natural Sciences and Engineering Research Council of Canada (RGPIN-2016-04292) and the Canadian Institute for Health Research (RN310864 – 375894).

## Author Contributions

Experiments were conducted by S.A.C., J.B., C.L.M., B.V. and T.L.S. and conceptualized by S.A.C. and B.G.H.; funding and supervision was provided by B.G.H.; the manuscript was written by S.A.C and B.G.H.

## Experimental Procedures

### Mouse Strains and Embryo Collection

All mice were held at the British Columbia Children’s Hospital Research Institute Animal Care Facility and ethical procedures were followed according to protocols approved by the University of British Columbia Animal Care Committee. All mice were maintained on a regular chow diet *ad libitum* and housed up to 5 mice per cage on a 12-hour light/dark cycle. Timed matings were used to determine embryonic stages and the morning of vaginal plug discovery was considered embryonic day 0.5 (E0.5). Previously generated *Dpy30*^flox/flox^ mice (Campbell et al., 2019) were crossed to *Neurog3*-Cre driver mice (Schonhoff et al., 2004) to obtain conditional deletion of *Dpy30* exon 4 in endocrine progenitors. In all studies, knockout mice (*Dpy30ΔN*; *Neurog3*-Cre; *Dpy30*^flox/flox^) were compared to Cre-negative littermate controls (*Dpy30*^flox/flox^ or *Dpy30*^flox/wt^). At noon on the day of the experimentally determined time point (e.g., E14.5), embryos were harvested by hysterectomy, dissected under an Olympus dissecting microscope in ice-cold phosphate buffered saline (PBS) and tail clippings were taken for genotyping. All knockout embryos were stained for DPY30 and/or H3K4 methylation to determine recombination efficiency prior to further analysis. Embryos with insufficient loss of DPY30 or H3K4 methylation (< 80%) were not studied. For all experiments, sex-specific differences were not anticipated and embryo sex was not determined.

### Pancreas and Islet Isolations

Mice were anesthetized with isoflurane inhalation and checked for toe pinch reflexes prior to euthanasia by cervical dislocation. To open the chest cavity, two lateral incisions were made in the abdomen away from the midline, followed by medial incisions up and down the midline and through the ribcage and diaphragm. The pancreas was dissected by cutting all connections to the spleen, stomach, intestine and liver before placing the tissue directly into 4% PFA in PBS on ice, followed by fixation overnight at 4°C.

For islet isolations, mice were euthanized by decapitation and blood was drained. The common bile duct was clamped at the duodenum and a 30G needle was used for perfusion of 3 mL Collagenase XI (1000 U/mL, Sigma-Aldrich) in Hank’s Balanced Salt Solution (HBSS) into the common bile duct. The pancreas was dissected and further enzymatically digested in an additional 2 mL Collagenase XI for 15 minutes in a 40°C water bath before 3 minutes of mechanical shaking. Pancreas tissue was washed twice with 25 mL HBSS with 1 mM CaCl_2_ before filtering through a 40 or 70 μm cell strainer (depending on mouse age). Islets collected on the strainer were inverted and rinsed into a petri dish with RPMI 1640 media (11 mM D-glucose) supplemented with 10% FBS, 50 U/mL penicillin/streptomycin and 2 mM L-glutamine, and either recovered overnight at 37°C in a 5% CO_2_ humidified incubator or immediately handpicked for experiments under a dissecting microscope.

### Tissue Processing and Histology

Embryonic and pancreatic tissue was fixed at 4°C in 4% PFA overnight (whole embryos) or for 5 hours (dissected stomach, pancreas, spleen and intestine ≥ E15.5). All tissue was processed through a series of graded ethanol and xylene de-hydration steps before embedding in paraffin wax. Briefly, fixed embryos and tissues were washed 3x 10 minutes in PBS and processed in cassettes through 50% ethanol (embryos only), 70% ethanol, 2x 30 minutes of 95% and 100% ethanol, 2x 30 minutes of xylene, followed by 2x 1 hour in melted paraffin. Embryonic or pancreatic tissue was cut into 5 μm sagittal sections using a Microtome and mounted onto Superfrost Plus slides.

### Immunostaining, Imaging and Analysis

Paraffin slides were processed through graded xylene and ethanol re-hydration steps, followed by a 10 minute heat-induced antigen retrieval (10 mM sodium citrate, pH 6.0) at 95°C and 1 hour blocking (5% FBS in PBS) at room temperature. Briefly, slides were processed through 3x 5 minutes xylene, 2x 5 minutes 100% ethanol, 5 minutes 95% ethanol, 5 minutes 70% ethanol and 10 minutes in PBS on a shaker prior to antigen retrieval. Slides were cooled for 5 minutes under cold running tap water and washed for 5 minutes each in de-ionized water and PBS on a shaker before circumscribing with a Super PAP pen (Thermo Fisher Scientific) and blocking. Primary antibodies were incubated at 4°C overnight with dilutions in blocking solution (see Table below for antibody information). The following day, slides were washed 3x 10 minutes in PBS and incubated with secondary antibodies in PBS for 1 hour at room temperature in a humidified dark chamber. Finally, slides were washed 3x 10 minutes in PBS before mounting with Prolong Gold mounting solution. Slides were imaged on a Leica TCS SP8 Confocal microscope or tiled on an Olympus Bx61 microscope and analyzed using CellProfiler software with custom pipelines.

| **Antibody** | **Host Species** | **Manufacturer** | **Catalog #** | **Dilution** |
| --- | --- | --- | --- | --- |
| Chromogranin A | Mouse | Santa Cruz | sc-393941 | 1:100 |
| DPY30 | Rabbit | Abcam | ab214010 | 1:500 |
| DPY30 | Rabbit | Bethyl | A304-296A | 1:10,000 |
| DPY30 | Rabbit | Atlas Antibodies | HPA043761 | 1:1000 |
| Gastrin | Goat | Santa Cruz | sc-7783 | 1:50 |
| Glucagon | Rabbit | Santa Cruz | sc-13091 | 1:500 |
| Glucagon | Mouse | Sigma-Aldrich | G6254 | 1:2000 |
| H3K4me1 | Rabbit | Abcam | ab8895 | 1:1000 |
| H3K4me3 | Rabbit | Abcam | ab8580 | 1:1000 |
| H3K4me3 | Rabbit | Cell Signaling | C42D8 | 1:1000 |
| Insulin | Guinea Pig | Abcam | ab7842 | 1:1000 |
| Insulin | Guinea Pig | Agilent | IR-002 | 1:4 |
| NEUROG3 | Mouse | DSHB | F25A1B3-c | 1:100 |
| Somatostatin | Goat | Santa Cruz | sc-7819 | 1:1000 |
| SOX9 | Rabbit | Millipore | AB5535 | 1:2000 |
| DAPI |  | Thermo Fisher Scientific | D9542 | 1:5000 |
| TOPRO-3 |  | Thermo Fisher Scientific | T3605 | 1:5000 |
| Alexa Fluor 488 anti-Rabbit | Donkey | Thermo Fisher Scientific | A-21206 | 1:500 |
| Alexa Fluor 555 anti-Guinea Pig | Goat | Thermo Fisher Scientific | A-21435 | 1:500 |
| Alexa Fluor 546 anti-Mouse | Goat | Thermo Fisher Scientific | A-11003 | 1:500 |
| Alexa Fluor 488 anti-Goat | Donkey | Thermo Fisher Scientific | A-11055 | 1:500 |
| Alexa Fluor 647 anti-Mouse | Donkey | Thermo Fisher Scientific | A-31571 | 1:500 |
| Alexa Fluor 594 anti-Rabbit | Donkey | Thermo Fisher Scientific | A-21207 | 1:500 |

### Morphometric and Intensity analysis

Cell quantifications were determined by taking serial sections at set intervals throughout the entire embryonic (every 30-60 μm) and postnatal (every 90 μm) pancreas. At least 10 images per replicate were captured on a Leica TCS SP8 Confocal microscope. Images were analyzed using CellProfiler software to quantify relative cell fractions or H3K4me3 immunostaining intensity. The number of NEUROG3^+^ nuclei was determined manually. Total CHGA^+^ area in the P24 pancreas was determined from staining islets for CHGA on serial sections through the entire pancreas spaced 90 μm apart. Images were tiled on an Olympus Bx61 microscope and analyzed relative to total pancreatic area using CellProfiler.

### Western Blot Analysis

Islets were lysed for 5 minutes at 95°C in Laemmli sample buffer (2% SDS, 10% glycerol, 60 mM Tris pH 6.8) containing 1 mM NaF, 1 mM PMSF and protease inhibitor cocktail (Roche). The Pierce BCA kit (Thermo Fisher Scientific) was used to determine lysate total protein concentrations and 5 μg protein was re-boiled for 5 minutes with 5% β-mercaptoethanol. Proteins were separated on 4-15% acrylamide gels and transferred for 60 minutes at 12.0 V to PVDF membranes (Millipore). Membranes were blocked in 5% BSA in TBS (20 mM Tris pH 7.6, 150 mM NaCl) containing 0.1% Tween-20 (TBS-T) for 1 hour at room temperature before overnight incubation at 4°C with primary antibodies in 5% BSA/TBS-T. Primary antibodies were as follows: mouse anti-actin (DSHB, catalog # JLA20-c, 1:2500) and rabbit anti-H3K4me3 (CST, catalog # 9751, 1:1000). Membranes were washed 3x 10 minutes in TBS-T before incubated in secondary antibodies for 1 hour at room temperature. Secondary antibodies were as follows: HRP-conjugated anti-rabbit IgG (CST, catalog # 7074, 1:10,000) and HRP-conjugated anti-mouse IgG (Jackson Immunoresearch, catalog # 115-035-174 1:10,000). Membranes were washed 3x 10 minutes in TBS-T and signals were detected using ECL reagent and radiographic film. Antibodies were then stripped in mild stripping buffer (200 mM glycine 0.1% SDS, 1% Tween-20, pH 2.2) for 20 minutes at room temperature and washed 4x 10 minutes in TBS-T. The membrane was re-blocked and probed with rabbit anti-H3K4me1 (Abcam, catalog # 8895, 1:100,000), then re-stripped, blocked and probed with rabbit anti-H3 (Abcam, catalog # ab1791, 1:250,000). The resulting signals were quantified using ImageJ.

### Blood Glucose and Islet Insulin Measurements

Mice were monitored after weaning at P21 for body mass and random blood glucose by a OneTouch® Ultra® 2 glucometer. For intraperitoneal glucose tolerance tests (IPGTTs), mice were fasted for 6 hours prior to IP injection of 2 g/kg 20% D-glucose (Sigma-Aldrich) in water with a 26G needle. Blood glucose measurements were obtained from the tail vein prior to injection (T_0_) and 15, 30, 60 and 120 minutes after injection. For islet insulin content, isolated islets were lysed in RIPA buffer (25 mM Tris pH 7.6, 150 mM NaCl, 1% NP40, 1% sodium deoxycholate and 0.1% SDS) for 5 minutes at 95°C. Islet insulin content was determined from the lysate by the Mouse Ultrasensitive Insulin ELISA kit (ALPCO) and a Spectramax 190 plate reader (Molecular Devices). Insulin measurements were normalized to islet lysate DNA concentration determined by the Qubit dsDNA HS Assay (Thermo Fisher Scientific).

### RNA Extraction and qPCR Analysis

RNA was extracted from lysed cells by pipetting in TRIzol reagent (Thermo Fisher Scientific) and combined with 1/5 the total volume of chloroform. Samples were inverted 10x before centrifugation at 12,000xg for 15 minutes at 4°C. The aqueous layer was mixed with an equal volume of ice-cold 70% ethanol and further processed with the PureLink RNA Mini Kit (Ambion). Complementary DNA (cDNA) was generated using SuperScript III Reverse Transcriptase (Thermo Fisher Scientific) and qPCR experiments were carried out using 0.25-2 μL of cDNA per reaction. Both Fast SYBR Green and TaqMan chemistry (Thermo Fisher Scientific) was used with purchased or custom designed primers (Primer3) to detect exon-intron boundaries. Samples were loaded onto 384-well plates in triplicate and run on a ViiA 7 Real-Time PCR system (Thermo Fisher Scientific) where gene expression was normalized to β-actin and determined using the ΔΔC_t_ formula.

### RNA-Sequencing Analysis

Islets from 3 control and 3 *Dpy30ΔN* male mice were isolated at P24 and 100-200 islets per mouse were handpicked into 1 mL of TRIzol reagent (Thermo Fisher Scientific). Extracted RNA was treated with TURBO DNase I (Thermo Fisher Scientific) for 20 minutes at 37°C to remove genomic DNA. For each sample at P24, mRNA was enriched from 1000 ng of total RNA using the NEBNext Poly(A) mRNA Magnetic Isolation Module (NEB) and 6 cDNA libraries were prepared with the NEBNext Ultra II Directional RNA Library Prep Kit for Illumina (NEB) with 9 amplification cycles. Indexed libraries were analyzed for size distribution using the Agilent Bioanalyzer High Sensitivity DNA chip and quantified using the Qubit dsDNA HS Assay (Thermo Fisher Scientific). Pooled libraries were sequenced on an Illumina NextSeq500 platform for 2 × 43 nucleotide paired-end reads.

For analysis of islet RNA-seq libraries, the resulting FastQ sequencing files were combined, filtered (phred33) and trimmed to remove adapters with Trimmomatic (Bolger et al., 2014). Salmon (Patro et al., 2017) was used to quasi-map, assemble and quantify transcripts using the mouse GRCm38/\VM24 transcriptome. Differential gene expression analysis was performed using DEseq2 (Love et al., 2014) in R, with the apeglm algorithm used to shrink log fold changes (Zhu et al., 2019). Gene set enrichment analysis was performed with g:Profiler (Raudvere et al., 2019).

Previously published RNA-seq data from E13.5 *Neurog3*^HI^ endocrine progenitors and E17.5 INS1+ pancreatic β-cells (Yu et al., 2018) was combined with RNA-seq data from P1, P7, P14, P21 and P28 pancreatic islets (Zeng et al., 2017) and was processed as described above. The transcripts per million were compiled to create a developmental timeline.

### ChIP-Sequencing Analysis

We used published ChIP-seq data (H3K4me3, H3K4me1, H3K27ac and H3K27me3) from E13.5 *Neurog3*^HI^ and adult islets (Lu et al., 2018; Tennant et al., 2013) and previously published transcription factor binding data for FOXA2, MAFA, PDX1 and NEUROD1 (Tennant et al., 2013), and for NKX2-2 (Gutiérrez et al., 2017), NKX6-1 (Taylor et al., 2013), PAX6 (Swisa et al., 2017), RFX6 (Piccand et al., 2014). All data were mapped with bowtie2 to the GRCm38/mm10 genome and peaks for the transcription factors called with MACS2. We used Repitools to plot the distribution of each histone mark at the transcriptional start sites (TSS, ± 3 kb) of the differentially expressed gene sets. We used ChromHMM (Ernst and Kellis, 2012) using a *k*-value of 10 to characterize the chromatin states of *Neurog3*^HI^ and islet cells. We determined the percent coverage of each state at the TSS (± 2 kb) of the differentially expressed gene sets using the coverage function of bedtools2 (version 2.29.2). Based on the hierarchy of chromatin states and highest percent coverage, each gene was defined a chromatin state from the *Neurog3*^HI^ and islet data sets.

### Statistics

Data are expressed as mean ± standard deviation (SD) unless otherwise specified and all experiments were carried out at minimum in triplicate. Statistical analyses were performed using GraphPad Prism 8 Software. Statistical significance was determined using unpaired, two-tailed Student’s t-tests for comparisons between two groups, and two-way ANOVA with Sidak’s multiple comparisons post-hoc tests were used for repeated measures comparisons between more than two groups (unless otherwise specified), with * indicating *P* < 0.05, ** indicating *P* < 0.01, *** indicating *P* < 0.001 and **** indicating *P* < 0.0001.

