## Supplementary Figures 1-3 for "H3K4 trimethylation is required for postnatal pancreatic endocrine cell functional maturation"

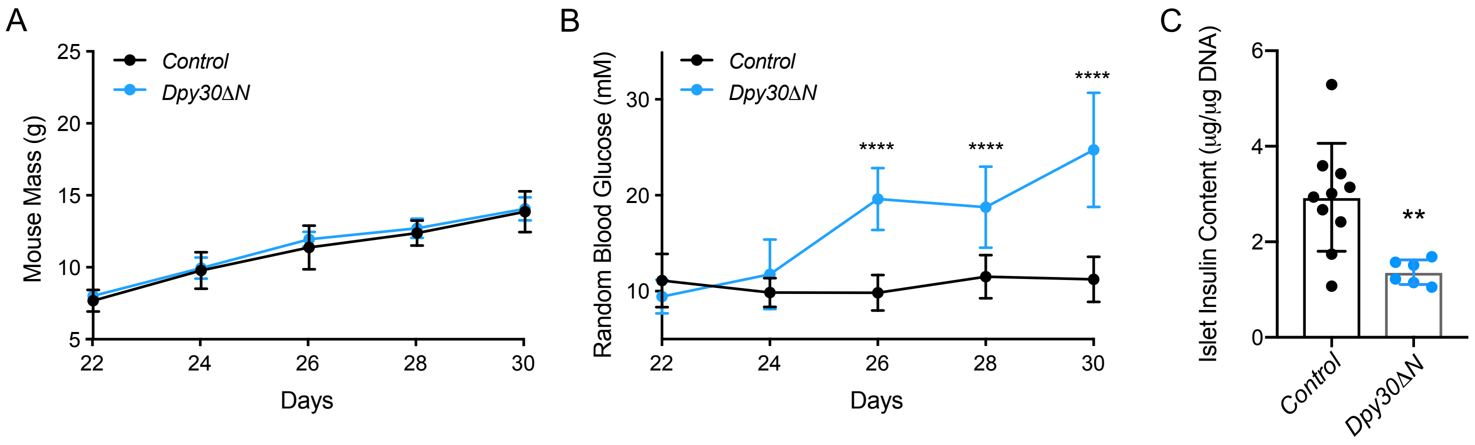


**Supplemental Figure 1:** Female *Dpy30ΔN* mice develop hyperglycemia and have islets with reduced insulin content. **(A)** Mouse body mass measurements from female control and *Dpy30ΔN* animals. **(B)** Random blood glucose measurements from female *Dpy30ΔN* and control mice between P22 and P30. **(C)** Total insulin content from P24 female control and *Dpy30ΔN* islets. Data are presented as mean ± SD; n = 6; multiple t-tests with Holm-Sidak correction for multiple comparisons.

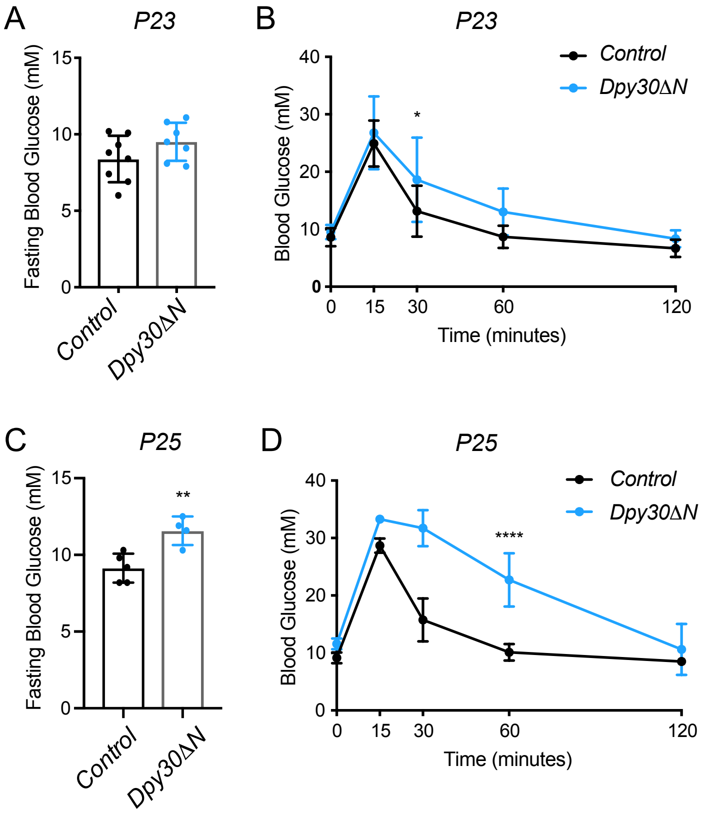


**Supplemental Figure 2:** Male *Dpy30ΔN* mice develop hyperglycemia and impaired glucose tolerance. **(A)** Blood glucose measurements after a 6 hour fast in P23 control and *Dpy30ΔN* mice. **(B)** IPGTT of 2 g per kg body mass IP glucose in P23 control and *Dpy30ΔN* mice following a 6 hour fast. **(C)** Blood glucose measurements after a 6 hour fast in P25 control and *Dpy30ΔN* mice. **(D)** IPGTT of 2 g per kg body mass IP glucose in P25 male control and *Dpy30ΔN* mice following a 6 hour fast. Data are presented as mean ± SD; n= 6; unpaired, two-tailed Student's t-test; repeated measures two-way ANOVA with Sidak's multiple comparison post-hoc test.

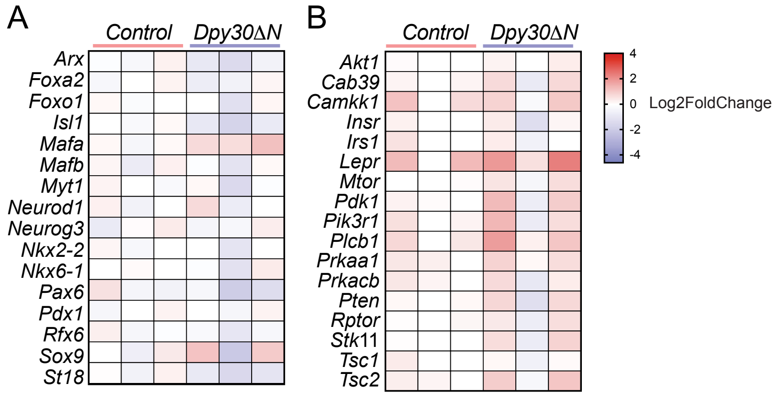


**Supplemental Figure 3:** Expression of key β-cell transcription factors and mTOR signalling pathway genes is largely unaltered in *Dpy30ΔN* mice.
**(A)** Heatmap of select transcription factor gene expression (log2 fold change) comparing 3 biological replicates across P24 control and *Dpy30ΔN* islets. **(B)** Heatmap of select mTOR pathway gene expression (log2 fold change) comparing 3 biological replicates across P24 control and *Dpy30ΔN* islets.
